# Prediction of gene cluster function based on transcriptional regulatory networks uncovers a novel locus required for desferrioxamine B biosynthesis

**DOI:** 10.1101/2024.06.10.598258

**Authors:** Hannah E. Augustijn, Zachary L. Reitz, Le Zhang, Jeanine A. Boot, Somayah S. Elsayed, Gregory L. Challis, Marnix H. Medema, Gilles P. van Wezel

## Abstract

Bacteria produce a plethora of natural products that are in clinical, agricultural and biotechnological use. Genome mining revealed millions of biosynthetic gene clusters (BGCs) that encode their biosynthesis, and the major challenge is to predict the bioactivities of the molecules these BGCs specify, and how to elicit their expression. Here, we present an innovative strategy whereby we harness the power of regulatory networks combined with global gene expression patterns to predict BGC functions. Studying the regulon of iron master regulator DmdR1 in *Streptomyces coelicolor* combined with co-expression data and large-scale comparative genome analysis identified the novel *desJGH* gene cluster. Mutational and metabolomics analysis showed that *desJGH* is required for biosynthesis of the clinical drug desferrioxamine B. DesJGH thereby dictate the balance between the structurally distinct desferrioxamines B and E. We propose regulation-based genome mining as a promising approach to functionally prioritize BGCs to accelerate the discovery of novel bioactive molecules.

## INTRODUCTION

Within the genetic blueprint of microorganisms lies an immense reservoir of chemical potential, which likely constitutes the mechanistic basis for numerous microbiome-associated phenotypes and offers a rich source of raw materials for discovery and development of among others antibiotics, anticancer agents, immunosuppressants, crop protection agents, and industrial ingredients ^1,2^. Genome mining efforts have led to the identification of millions of biosynthetic gene clusters (BGCs) predicted to encode the biosynthesis of many thousands of natural product scaffolds ^3^. However, only an estimated 3% of these specialized metabolites have undergone experimental characterization thus far, leaving a vast amount of untapped chemical diversity yet to be explored ^4^.

Identifying the diverse roles of specialized metabolites in microbiome interactions is highly challenging, primarily due to the dynamic nature of the host environment and the difficulties in replicating such conditions in laboratory settings. Moreover, while these molecules exhibit a wide range of functions, only a small fraction of metabolites will directly contribute towards microbiome-associated phenotypes such as disease suppression or growth promotion, or have the necessary properties to yield the next generation of crop protection agents, antibiotics, or food additives ^5–7^. As a result, there is a pressing need for generalized strategies to predict the functions of specialized metabolites, enabling us to understand their mechanistic roles in inter-organismal interactions and to gauge their usefulness for industrial and clinical applications.

A major aim in current natural product discovery is to identify ways to reduce the genetic space of sequenced BGCs to manageable numbers, to inform scientists on which BGCs to prioritize in the search for novel bioactivity. Historically, scientists have investigated two dimensions, namely the molecular space via high-throughput screening of compound and strain libraries, followed by the genomic space in the 21^st^ century, by investigating BGCs in sequenced genomes, based on the identification of enzyme-coding genes ^8^. Perhaps the most advanced strategy for the latter has thus far been target-based genome mining, which uses self-resistance genes inside BGCs as beacons for recognizing the macromolecular targets of their products. However, the presence of recognizable self-resistance genes seems to be limited to a mere 5-10% of BGCs, necessitating complementary methods to predict the functions of the remaining specialized metabolic diversity ^9,10^.

We anticipate that an attractive alternative would be regulation-guided approaches, given that the regulatory system plays a pivotal role in the transcription of BGCs. Overexpression or inactivation of cluster-situated regulatory genes have been used to activate their expression ^11–13^. For example, targeting BGCs containing *Streptomyces* antibiotic regulatory protein (SARP) family regulators enabled the discovery of novel antibiotic BGCs ^14,15^. Also, the Identification of Natural compound Biosynthesis pathways by Exploiting Knowledge of Transcriptional regulation (INBEKT) strategy was able to unveil a previously undetectable BGC by identifying regulatory binding sites of the zinc-dependent regulator ZuR ^16^. These early successes at the single-gene or single-BGC level indicate that genome-wide analysis of regulatory networks may be even more successful at unveiling BGC functions.

Here, we introduce a computational omics strategy that leverages genome-wide gene regulation information to provide functional predictions of BGCs in microbes. This novel approach connects genome-wide regulatory information derived from transcription factor binding site (TFBS) prediction to gene co-expression networks, thereby associating genes to functions. Genome-wide regulatory analysis of BGCs of *Streptomyces coelicolor* M145 in combination with co-expression patterns unveiled a novel BGC that had escaped detection by current genome mining software tools. Subsequent mutational analysis and metabolic profiling experiments showed that this BGC plays an important role in the biosynthesis of the well-studied siderophore desferrioxamine B. These results illustrate the potential of our method to infer BGC function, facilitate the detection and prioritization of novel BGCs and ultimately pave the way for identifying genes responsible for the biosynthesis of novel bioactive molecules.

## RESULTS AND DISCUSSION

### Identifying functional associations through gene regulatory networks

A major challenge in genome-mining-based drug discovery lies in prioritizing BGCs within the vast unexplored biosynthetic space, and in particular finding novel ways to predict their function. We hypothesized that regulatory networks that control BGC expression might form a new, third, dimension for screening for potential functions, complementing phenotypic and genomic screening. The concept we propose is that if an unknown BGC (or any cluster of genes) is predicted to be controlled by a transcriptional regulator that responds to a known signal and is connected to a specific physiological response, that BGC may functionally relate to known BGCs controlled in a similar manner.

To develop such a regulation-based genome mining strategy and assess its validity, we chose to focus on the model organism *Streptomyces coelicolor* M145. This microbe, belonging to the phylum Actinomycetota, is renowned for its exceptional ability to produce a wide array of bioactive compounds, making it an interesting target for natural product discovery ^17–20^. Moreover, it is the bacterial species with currently the largest number of functionally characterized BGCs, with 17 out of its 27 BGCs having been connected to the production of a known metabolite, making it an ideal organism to assess how well regulation connects to function ^21^. To investigate the functional relationships between this microbe’s regulatory machinery and specialized metabolite biosynthesis, we investigated the binding of transcription factors (TFs) to their corresponding binding sites (TFBSs). For this purpose, we used the regulatory data of the LogoMotif database ^22^. Seventeen precalculated and manually curated position weight matrices (PWMs) associated with TFs in this database were used for genome-wide predictions of 730 TFBSs, using automated computational matching. Based on these predictions, a gene regulatory network (GRN) was constructed in which TFBSs were identified within BGC regions predicted by antiSMASH (Fig. 1a). A total of 81 TFBSs were found within antiSMASH BGC regions; 55 of these were at the region peripheries and putatively unrelated to specialized metabolite biosynthesis. To identify which TFBSs were truly linked to biosynthetic pathways, we then refined the boundaries of the BGCs (Table S1) using literature evidence and gene co-expression patterns (see below). This resulted in the identification of 17 low-confidence and 9 medium/high-confidence BGC-TFBS associations each matching the physiological or ecological functions associated with the corresponding regulon (Fig. 1a). These findings agree with existing experimental analyses, thus reinforcing the utility of our approach in accurately identifying BGC-TFBS connections (Fig. 1b). For example, there is a clear correlation between TFBSs of the zinc uptake regulator (Zur) and the zinc-regulated coelibactin locus^23^, as well as between the pleiotropic antibiotic biosynthesis regulator AfsQ1 and the antibiotic coelimycin P1 ^24^. Additionally, we observed a connection between the iron-dependent regulator DmdR1 and the biosynthesis of two iron-chelating compound families that function as siderophores: the desferrioxamines (DFOs) and coelichelin ^25,26^.

**Figure 1.**
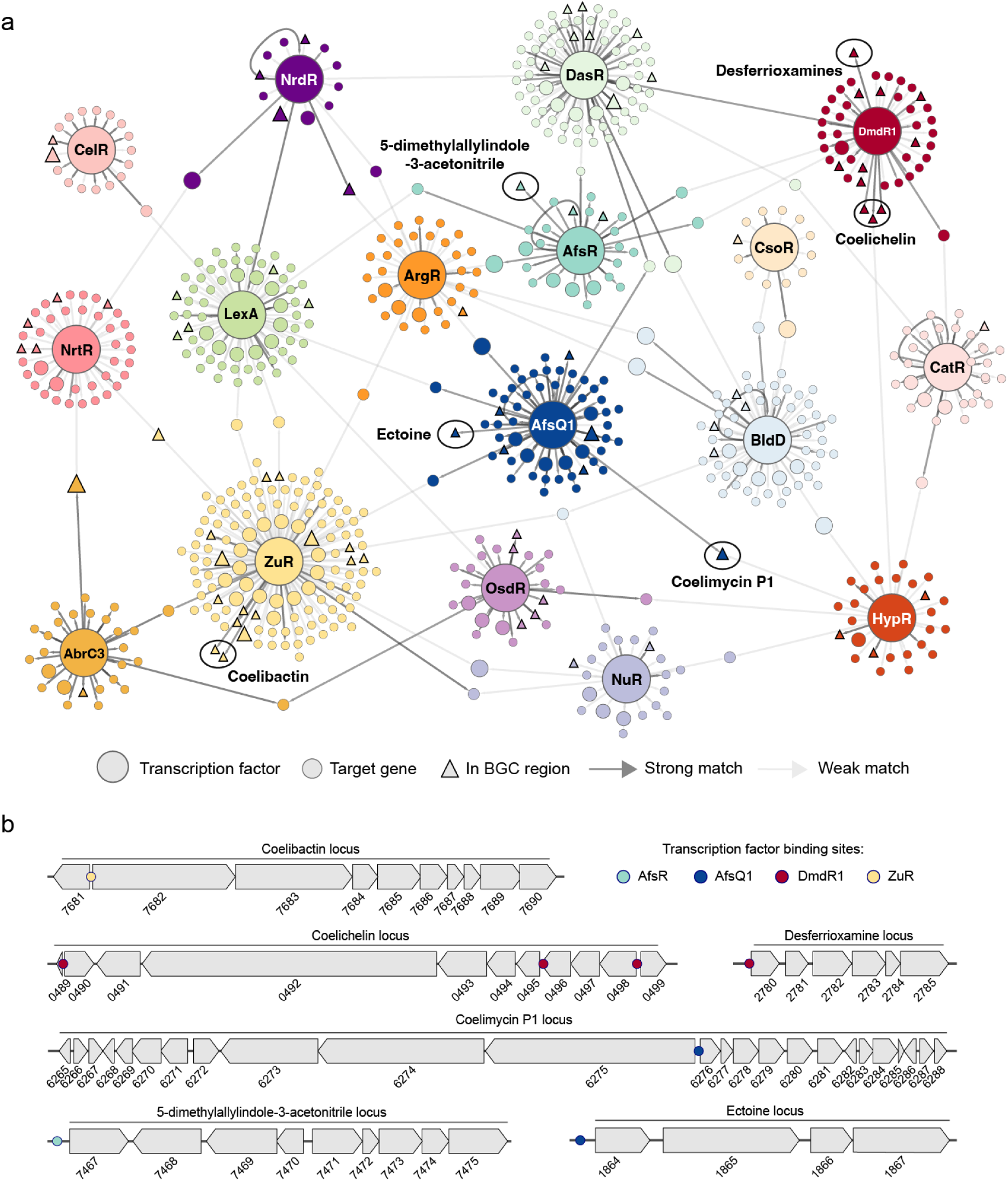
**a**, Predicted gene regulatory network of *Streptomyces coelicolor* based on 17 well-known regulators. Each node in the network represents a (regulatory) gene, and every edge represents a regulatory interaction between two nodes. The edges colored in dark gray indicate strong PWM prediction scores, while the lighter gray shades represent weaker interactions. Matches within BGC regions are depicted as triangles. In six regions (black circled), the matches fall within a co-expressed region, highlighting their functional relation to these compounds. **b,** Representation of the four co-expressed regions, including the locations of their detected TFBSs as colored dots. All predicted TFBSs have been experimentally validated in pre-existing work.

### Co-expression analysis and operon-level expansion of the predicted DmdR1 regulon

Next, we aimed to go beyond antiSMASH-detectable BGCs and assess if we could infer the function of any uncharacterized operons and gene clusters using regulatory predictions. Expectedly, the predicted DmdR1 regulon exhibited a clear functional association with siderophores, as evidenced by the connection between its binding sites and known siderophore BGCs ^27^. Therefore, we focused on exploring the functional connection between DmdR1 binding sites (*iron boxes*) and iron metabolic genes. A critical issue when using PWMs is the large number of false positive TFBS hits. To address this, we refined the general LogoMotif detection threshold for DmdR1 to be more accurate for *S. coelicolor* by applying the principles previously described for the calibration of the PREDetector algorithm ^28^. This approach involves an analysis of the distribution of hits and the ratio of hits in non-coding versus coding regions (Fig. S1). The results demonstrated that higher PWM match scores correlated with a greater frequency of hits detected in non-coding regions, where iron boxes are typically found. By calculating the median score of the non-coding to coding ratio, we established a refined threshold of 22.875, leading to the identification of a total of 39 predicted DmdR1 binding sites (Table S2). Among these 39 predicted binding sites, we identified 25 unique binding site locations, 22 of which corresponded to previously reported DmdR1 target genes. Based on these predictions, we identified three novel putative DmdR1 target genes: SCO2114, SCO2275, and SCO5998.

Bacterial regulons consist not only of genes with TFBSs in their regulatory region, but also any downstream co-operonic genes. DmdR1-controlled operons were predicted using a co-expression analysis of a previously published transcriptome. The RNA-Seq dataset of Lee *et al*.^29^ was chosen for its relatively high sample count (22 for *S. coelicolor*) and the study’s focus on iron restriction. Reads were retrieved from NCBI SRA and mapped to the *S. coelicolor* M145 genome, and gene count data were processed using previously reported techniques to generate a pairwise gene co-expression matrix ^30,31^. Of the 30 predicted DmdR1 target genes with a significant PWM match score, 26 were anti-correlated with transcription of *dmdR1* (Pearson correlation coefficient [PCC] < –0.43, *p*<0.05, Fig. 2a), including newly predicted target genes SCO2114, SCO2275, and SCO5998. The co-expression data support the minimum PWM match score of 22.875; below this threshold, no mean anti-correlated expression was identified. Only a single gene with a significant PWM score, the GntR-type regulator SCO6159, was positively co-expressed with *dmdR1* (PCC = 0.69), and the transcription pattern of three putative target genes did not correlate significantly with that of *dmdR1*, suggesting a more complicated regulation by multiple transcription factors. DmdR1 target genes were placed into predicted operons using the gene co-expression matrix, as well as strand and intergenic distance, expanding the putative direct regulon of DmdR1 from 25 to 58 genes, which are found across 16 genomic loci (Fig. 2b). A description of the predicted DmdR1 regulon, including functional predictions, is presented in SI Discussion 1. As expected, DmdR1 binding sites were recovered in the coelichelin and desferrioxamine BGCs but not the ZuR-controlled coelibactin BGC, supporting the use of regulatory analysis for linking metallophore BGCs to their corresponding metal. Other logical gene annotations present in the regulon include siderophore-independent iron acquisition, mobilization of stored iron, and oxidative stress response.

**Figure 2.**
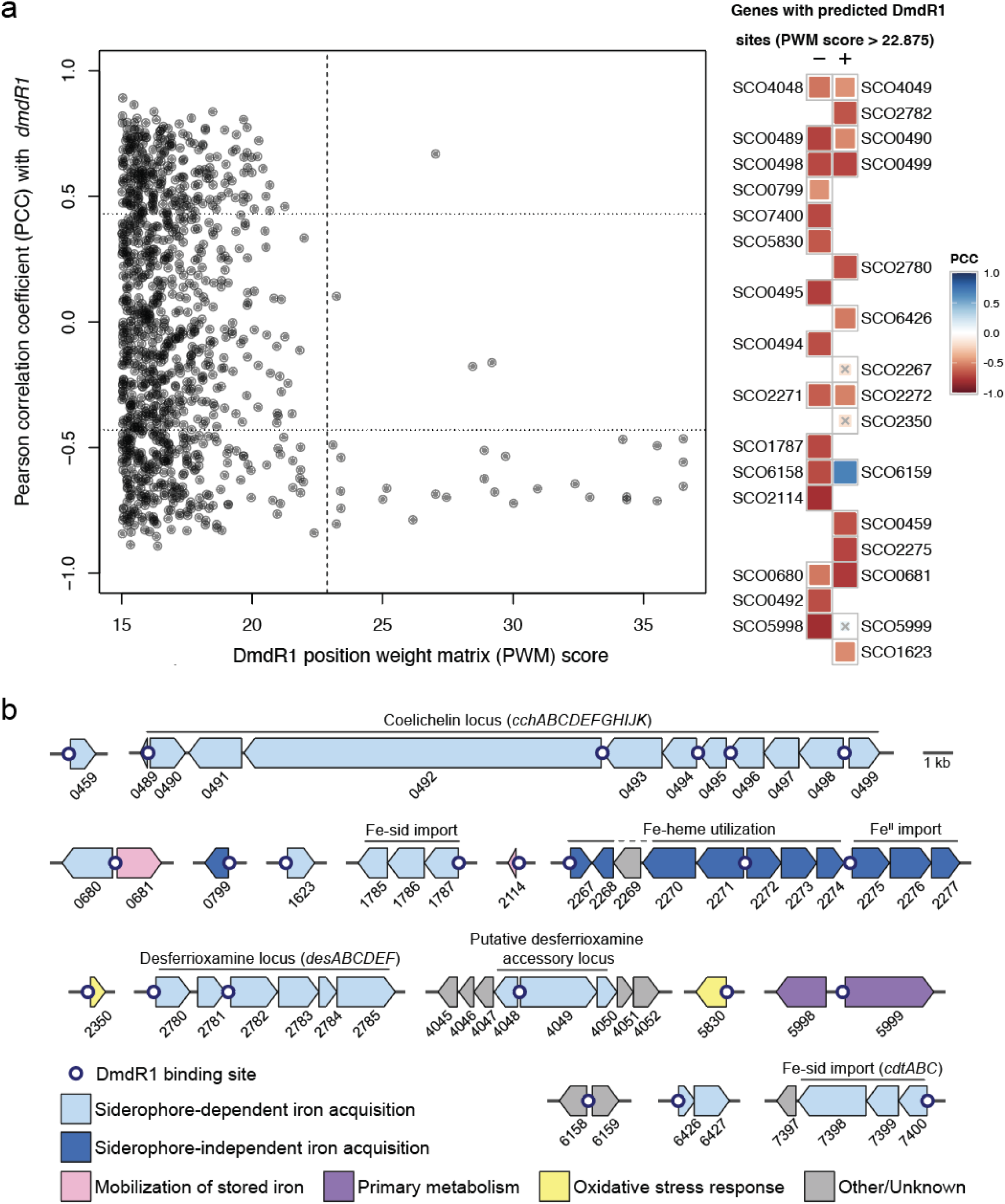
a, Anti-correlation of gene expression between *dmdR1* and its predicted regulon. Left: Pearson correlation coefficients (PCCs) between *dmdR1* and all genes with a DmdR1 position weight matrix (PWM) score greater than 15 in their regulatory region. The vertical dashed line marks the refined PWM score threshold of 22.875. The horizontal dotted lines mark PCC = ±0.43, corresponding to an adjusted *p*-value of 0.05. Right: Target genes immediately downstream of a predicted DmdR1 binding site, ordered by decreasing PWM score. Plus and minus indicate the strand of the target gene. Genes marked with an × did not have significant co-expression with *dmdR1*. Binding site details are given in Table S2. **b,** The putative regulon of DmdR1 in *S. coelicolor* M145. White dots indicate predicted DmdR1 binding sites. Genes are labeled by SCO number and colored by putative function. Clusters are drawn to scale, and arrows represent the direction of transcription.

### Metabolic profiling of an unexplored DmdR1-controlled locus

This systematic mapping of the DmdR1 regulon then provided the opportunity to investigate whether new operons or gene clusters could be identified that would be predicted to function in iron acquisition. Upon close examination of all individual genes across the regulon, the uncharacterized region from SCO4045 to SCO4052 stood out due to sequence similarity to biosynthetic genes (Fig. 2b). Interestingly, SCO4050 encodes a protein similar to the *N*-acyltransferase DesC (encoded by SCO2784), which catalyzes the conversion of *N*-hydroxycadaverine to *N*-hydroxy-*N*-succinylcadaverine (HSC) and *N*-hydroxy-*N*-acetylcadaverine (HAC), the direct precursors of desferrioxamine B, *in vitro* ^32^. SCO4048 is a paralog of *desF* (SCO2781), which encodes ferrioxamine reductase. Furthermore, SCO4049 is homologous to genes designated as *desG* in other streptomycetes, and is predicted to encode a penicillin amidase family protein; phylogenetic analysis in Actinobacteria revealed that *desG*, if present, either colocalized with the DFO cluster, or with a separate DmdR1-controlled locus ^33^. DesG was originally hypothesized to increase DFO structural diversity by producing phenylacetic acid-capped derivatives in some strains; however, no arylated DFOs have been identified in *S. coelicolor*. Together, SCO4048, SCO4049, and SCO4050 (further referred to as *desJ*, *desG*, and *desH*, respectively) appear to comprise a previously undetected locus putatively related to DFO biosynthesis ^33^.

To analyze the role of the DmdR1-controlled locus in the production of DFOs, we applied the CRISPR-based editing system (CRISPR-BEST)^34^ to construct three knock-out mutants in which either SCO4048 (*desJ*), SCO4049 (*desG*) or SCO4050 (*desH*) had been inactivated. The system allows introduction of a premature stop codon in the target ORF, thus preventing the production of a functional protein. Using this method, we created null mutants of SCO4048 (*desJ*) with mutations W55* or Q68*, resulting in 186 aa or 173 aa truncation of the gene product, respectively. The introduction of a stop codon at W61 in SCO4049 (*desG*) led to a substantial 721 aa shortening, while mutations W43* or Q91* in SCO4050 (*desH*) resulted in truncations of 163 aa or 115 aa, respectively. PCR followed by DNA sequencing was used to verify the correctness of the knock-out mutants.

To obtain extracts for metabolomics, *S. coelicolor* M145 and its mutant derivatives were grown in a liquid iron-limited medium (ISP-2) for five days. The metabolites produced were adsorbed on Diaion® HP20 resin, which was subsequently extracted with methanol and analyzed using liquid chromatography-mass spectrometry (LC-MS), which revealed changes in the production of DFO-related metabolites in each of the mutants compared to the wild-type strain (Fig. 3a). The metabolites were annotated by matching the high-resolution mass spectrometry (HRMS) and tandem mass spectrometry (MS/MS) spectra to previously published ones (Fig. S2) ^35–37^. Statistical analyses showed that only the levels of desferrioxamine B (DFOB) were significantly increased in extracts of the *desJ* mutant as compared to the parental strain (Fig. S3). Metabolomic analysis of Δ*desG* and Δ*desH* revealed an approximate 1000-fold and 16-fold decrease in DFOB production, respectively (Fig. 3 and Fig. S3). Conversely, the mutants exhibited a significant increase in desferrioxamine E (DFOE) and its metal complexes, most likely as a result of the nearly abolished DFOB production.

**Figure 3.**
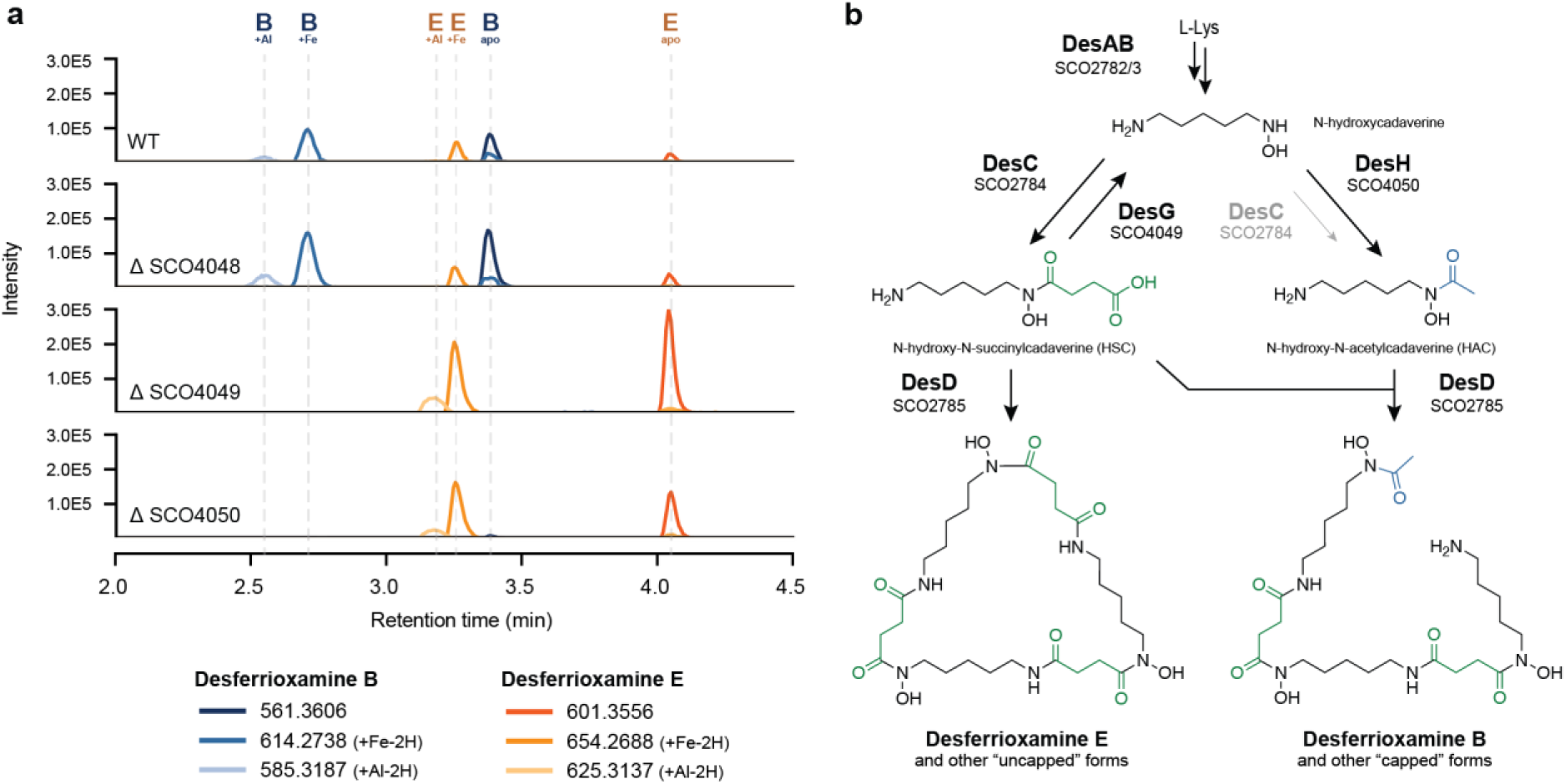
New model for biosynthesis of desferrioxamines B and E. a, Extracted ion chromatograms for *m/z* values corresponding to DFO-related metabolites in culture extracts of the knock-out mutants of SCO4048 (*desJ*), SCO4049 (*desG*) and SCO40450 (*desH*) compared to the parent *S. coelicolor* M145 strain. The *desG* mutant fails to produce DFOB, while a 16-fold decrease in DFOB biosynthesis was seen in *desH* mutants (c*f.* Fig. S3). **b,** Proposed biosynthetic pathway for assembly of desferrioxamines E and B. Main biosynthetic enzymes presented in bold face. DesG and DesH balance intracellular *N*-hydroxy-*N*-succinylcadaverine (HSC) and *N*-hydroxy-*N*-acetylcadaverine (HAC) concentrations by converting HSC to HAC. In the absence of DesG and/or DesH, the cells likely fail to produce sufficient levels of HAC, thereby strongly attenuating the production of DFOB. Although DesC has been shown to be able to catalyze the acetylation of *N*-hydroxycadaverine *in vitro*, the enzyme can only modestly compensate for the loss of DesH *in vivo*, underlining the important role played by DesG and DesH in DFOB production (Fig. S4).

We genetically complemented the mutants to determine if the effects were due solely to the gene inactivation and not to second-site mutations. For this, constructs were introduced that expressed the respective wild-type genes *desJ, desG* or *desH* from the constitutive *gap* promoter. The complementation constructs were based on vector pSET152 ^38^, which integrates at the bacteriophage ΦC31 attachment site on the *S. coelicolor* genome. The complemented mutants showed recovery of DFOB production in the complemented strains (Fig. S5). Taken together, our mutational analysis shows that the attenuation of DFOB production in the mutants can be fully explained by the inactivation of *desG* and *desH*.

DFOB and other capped desferrioxamines have been isolated from many *Streptomyces* strains, as well as several other Actinomycetota. To see if the proposed biosynthetic role for DesGH applies more generally to DFO biosynthesis in other Actinomycetota, we performed a meta-analysis of published DFO producers. In total, we identified reports of DFO production in 46 sequenced strains, comprising mostly *Streptomyces* species (n=34), as well as other Actinomycetota (n=7), Pseudomonadota (n=4), and one member of Bacteroidota (Table S3). Homologues of d*esG* and *desH* were found in 36 of the genomes, all Actinomycetota. One sequenced DFO producer, *Gordonia rubripertincta* CWB2, contained *desG* but not *desH*; however, the *G. rupripertincta* DFO locus is part of a larger BGC that putatively encodes the biosynthesis of the cryptic nocardichelins (see SI Discussion 2), and one of the two other acyltransferase genes in the BGC has presumably replaced *desH*. In all other cases, *desG* and *desH* are putatively co-operonic, and the two genes are fused in *Streptomyces atratus* and *Micrococcus* spp. CH3 and CH7. Among collected reports of DFO production, DFOB (Fig. 3b) and other acetyl, fatty-acyl, or aryl “capped” DFOs were common, isolated from 34 of 47 sequenced strains. However, in line with our discovery, the nine strains lacking *desGH* exclusively produced DFOE (Fig. 3b) and other “uncapped” DFOs with succinylated monomers (Fig. S6).

Based on the combination of the above data, we propose the following pathway for desferrioxamine biosynthesis in *S. coelicolor* (Fig. 3b). The biosynthesis of DFOE is encoded by the canonical biosynthetic locus *desABCD* (SCO2782-85): DesA and DesB convert L-lysine to *N*-hydroxycadaverine, DesC succinylates *N*-hydroxycadaverine to form HSC ^32^, and DesD cylcotrimerizes HSC to produce DFOE ^39^. In contrast, DesG (SCO4049) and DesH (SCO4050) enable DFOB production (Fig. 3). A recent study of DesD concluded that the relative intracellular concentrations of HSC and HAC must be controlled for DFOB formation ^39^. Previous investigations of DesC *in vitro* have shown that it is able to catalyze the conversion of *N*-hydroxycadaverine to both HSC and HAC, using succinyl and acetyl-CoA, respectively ^32^. However, the relative catalytic efficiency of these two processes has yet to be elucidated. Our experiments strongly suggest that the main function of DesC *in vivo* is to catalyze the production of HSC, while HAC results primarily from the action of DesH. We propose that DesG, which shows sequence similarity to amidases, de-succinylates HSC to regenerate *N*-hydroxycadaverine, which is then acetylated by the putative acetyltransferase DesH to boost the levels of HAC relative to HSC in high level DFOB producers. Gene fusions of *desGH* observed in some strains are equipped to exploit the high local effective concentration of *N*-hydroxycadaverine generated by the DesG domain, enabling the DesH domain to acetylate *N*-hydroxycadaverine before it can be re-succinylated. The production of DFOB in the Δ*desH* mutant is strongly attenuated but not abolished, consistent with the previously reported ability of DesC to catalyze acylation of *N*-hydroxycadaverine with acetyl-CoA in addition to succinyl-CoA (Fig. 3a). Taken together, these data indicate that DesC strongly prefers succinyl-CoA as a substrate over acetyl-CoA, and that DesG and DesH are required to ensure sufficient quantities of HAC are produced to support high level DFOB production *in vivo*. This biosynthetic model is in line with the available phylogenomic, metabolomic, and genetic evidence, as well as the canonical catalytic chemistry of DesG and DesH homologues.

## CONCLUSION

In conclusion, we have developed a novel computational omics strategy for functional inference of BGCs in microbes, which uses regulatory information to provide clues regarding their functional roles in inter-organismal interactions and to gauge their usefulness for industrial and clinical applications. Uniquely, this method leverages genome-wide gene regulation information derived from TFBS detection combined with gene co-expression network analysis to link biosynthetic genes to their potential functions. A key application of this method is showcased in our study of *Streptomyces coelicolor* M145, a well-studied model organism, where we predict the regulons of 17 well-known regulators and 9 high-confidence functional associations to known BGCs. Of these, we selected the iron-dependent repressor DmdR1 and its strong connection to the regulation of siderophore biosynthesis for showcasing the effectiveness of our approach. This analysis, which involved TFBS prediction of the DmdR1 regulon, alongside the detection of co-expression patterns under iron starvation conditions, allowed us to detect an uncharacterized gene cluster with a functional link to iron metabolism. Furthermore, we present evidence that the putative amidase and acyltransferase encoded by *desG* and *desH*, respectively, in this cluster collaborate in the efficient biosynthesis of desferrioxamine B by SCO4049 and SCO4050 CRISPR-cBEST knockout mutants and subsequent metabolic profiling experiments. These findings not only validated our hypothesis, but also enabled identification of a novel pathway within the complex biosynthetic route to desferrioxamines. Overall, our results demonstrate the effectiveness of our method in identifying and inferring the function of novel BGCs that escaped detection despite the availability of state-of-the-art genome mining tools. We anticipate that transcriptomics-guided regulatory genome mining, by combining function prediction with application of elicitors that may activate BGCs of interest, will provide pointers as to how to select and activate cryptic BGCs in the extant biosynthetic diversity. This will aid in the identification of their roles in microbiome interactions and guide the discovery of bioactive natural products that are of value for pharmaceutical, agricultural, and biotechnological applications.

## METHODS

### General

Default software parameters were used unless otherwise noted. Scripts are available at: https://github.com/zreitz/dmdR.

### Construction of the position weight matrix and sequence motif

Ten previously reported DmdR1 binding sites from *Streptomyces coelicolor* were collected from literature ^26^. Thereafter, the occurrences of each nucleotide across all positions of the sequences were counted to construct a position frequency matrix (PFM). This PFM was converted to a PWM by applying Bioconductor’s seqLogo v5.29.8 algorithm^40^, which calculates the log-likelihood of each nucleotide in the matrix, while taking into account the background nucleotide distributions. Additionally, the information content (IC) of the resulting PWM was calculated using Shannon’s entropy calculation methods. The IC was visualized as a sequence motif with the use of Logomaker v 0.8^41^.

### Identification of DmdR1 binding sites

The genome assembly of *Streptomyces coelicolor* A3(2) was downloaded from NCBI using accession GCA_000203835.1. The coding and non-coding regions, as well as the regions spanning from –350 bp to +50 bp relative to the start codons of each gene were extracted with MiniMotif^22^ (https://github.com/HAugustijn/MiniMotif). We employed MOODS v1.9.4.1^42^ to query these regions for DmdR1 PWM matches, using a p-value threshold of 0.01 and background distribution of 72% representing the GC percentage of *S. coelicolor*. The ratio of hits in non-coding versus coding regions was visualized using the R package ggplot2 ^43^.

### RNA-Seq data processing and co-expression analyses

*Streptomyces coelicolor* A3(2) RNA-Seq data, collected by Lee *et al*.,^29^ was retrieved from the European Nucleotide Archive (PRJEB25075).^44^ Raw read quality was assessed with FastQC.^45^ Reads were mapped to the reference genome NC_003888.3 using STAR v2.7.6a:^46^ Index files were generated with the parameters “--genomeSAindexNbases 10 –-sjdbGTFfeatureExon CDS”, and reads were aligned with the parameter “--alignIntronMax 1”. Mapped reads were indexed using SAMtools v1.3.1^47^ and visualized with the Integrative Genomics Viewer.^48^ Per-gene read count tables were generated with featureCounts v2.0.1^49^ using the parameters “-O –M –t CDS –s 2 –-fraction”.

The per-gene RNA-Seq count data was further analyzed in R. A minimum gene expression cutoff was applied (≥5 counts in 50% of samples), then counts were normalized by Trimmed Mean of M-values (TMM) and log_2_ transformed using a hyperbolic arcsine pseudocount ^50^. A co-expression bias associated with lowly– and highly-expressed genes (of unknown origin, but present in several other RNA-Seq datasets ^31^) was mitigated by regressing out the first principal component using the *sva_network* function from the *sva* package (Fig. S7)^30^. The resulting correlation matrix still had an expression-correlated broadening of correlation coefficients, which was corrected by spatial quantile normalization (Fig. S7)^31^ and used for further analyses. An all-to-all Pearson Correlation Coefficient (PCC) matrix with corrected two-sided Student p-values was calculated using the *corAndPValue* function from the package *WGCNA*.^51^ A p-value of 0.05 corresponded to a minimum absolute PCC value of 0.43. The correlation matrix was corrected for remaining expression-level-dependent PCC distribution broadening using spatial quantile normalization (*spqn::normalize_correlation*) with the following parameters: ngrp = 20, size_grp = 337, ref_grp = 18.^31^ Subsets of the resulting correlation matrix were used for all downstream analyses.

### Comparative genomics

Desferrioxamine core loci (*desABCD*) and accessory loci (*desGH*) were found in *Streptomyces* genomes using a modified version of antiSMASH 7 ^52^ (https://github.com/zreitz/antismash/tree/desGH-7-1). The “desABCD” rule requires matches to all of the following Pfam models with a maximum intergenic distance of 5 kbp: PF00282.22 (*desA*), PF13434.9 (*desB*), PF13523.9 (*desC*), and PF04183.5 (*desD*). The “desGH” rule requires matches to PF01804.21 (*desG*) and PF13523.9 (*desH*) with a maximum intergenic distance of 1 kbp. Genome assemblies for previously reported DFO producers (Table S3) were downloaded from NCBI Genbank on 21 Nov, 2023, in Genbank format using ncbi-genome-download^53^. The multiSMASH pipeline^54^ was used to scan the genomes with antiSMASH and tabulate the results ^52^. A gene phylogeny of the resulting desABCD loci was obtained from CORASON, run as part of BiG-SCAPE v1.1.5 using settings “--mix –-no-classify –-clans-off –-cutoffs 1” ^55^. The resulting phylogenetic tree was annotated using iTOL v5 ^56^.

### Bacterial strains and media

1. *E. coli* strains DH5ɑ and ET12567/pUZ8002^57^ were used for routine cloning and for interspecific conjugation, respectively. *E. coli* transformants were selected on Luria Bertani (LB) agar media containing the relevant antibiotics and grown O/N at 37 °C. *Streptomyces coelicolor* A3(2) M145 was used as parental strain to construct mutants. All media and routine *Streptomyces* techniques are described in the *Streptomyces* manual ^58^. Soy flour mannitol (SFM) agar plates were used to grow *Streptomyces* strains for preparing spore suspensions.

### Growth conditions and extraction

The cultures were grown in triplicate in 100 mL Erlenmeyer flasks with 1 g of Diaion® HP-20 resin (Resindion, Mitsubishi) in 15 mL of International *Streptomyces* Project-2 medium (ISP-2; yeast extract 4 g/L, malt extract 10 g/L and dextrose 4 g/L at pH 7.2). The medium was inoculated using 1 μL of spore stock and incubated in a rotary shaker at 30 °C. After five days of growth, the resin was vacuum filtered, washed three times with Milli-Q water, and extracted with 3 x 5 mL of methanol. The crude extracts were then dried, weighed, and dissolved in methanol at a final concentration of 1 mg/mL. Media blanks were extracted and prepared in a similar way as negative controls.

### LC-MS based metabolic profiling

Liquid chromatography-tandem mass spectrometry (LC-MS/MS) acquisition was performed using Shimadzu Nexera X2 ultra high-performance liquid chromatography (UPLC) system, with attached photodiode array detector (PDA), coupled to Shimadzu 9030 QTOF mass spectrometer, equipped with a standard electrospray ionization (ESI) source unit, in which a calibrant delivery system (CDS) is installed. A total of 2 µL of dissolved extracts were injected into a Waters Acquity HSS C18 column (1.8 μm, 100 Å, 2.1 × 100 mm). The column was maintained at 30 °C, and run at a flow rate of 0.5 mL/min, using 0.1% formic acid in H_2_O as solvent A, and 0.1% formic acid in acetonitrile as solvent B. A gradient was employed for chromatographic separation starting at 5% B for 1 min, then 5–85% B for 9 min, 85–100% B for 1 min, and finally held at 100% B for 3 min. The column was re-equilibrated to 5% B for 3 min before the next run was started. The LC flow was switched to the waste the first 0.5 min, then to the MS for 13.5 min, then back to the waste to the end of the run.

The MS system was tuned using standard NaI solution (Shimadzu). The same solution was used to calibrate the system before starting. Additionally, a calibrant solution made from ESI tuning mix (Sigma-Aldrich) was introduced through the CDS system, the first 0.5 min of each run, and the masses detected were used for post-run mass correction for the file, ensuring stable accurate mass measurements.

System suitability was checked by regularly measuring a standard sample made of the following compounds:

| compound | concentration (μg/mL) | retention time (min) | expected m/z |
| --- | --- | --- | --- |
| paracetamol | 25 | 2,375 | 152,0712 |
| caffeine | 5 | 3,246 | 195,0882 |
| prednisolone | 2,5 | 5,290 | 361,2015 |
| reserpine | 1,25 | 6,186 | 609,2812 |
| clomipramine | 1,25 | 6,379 | 315,1628 |

All the samples were analyzed in positive polarity, using data dependent acquisition mode. In this regard, full scan MS spectra (*m/z* 100–1700, scan rate 10 Hz, ID enabled) were followed by two data dependent MS/MS spectra (*m/z* 100–1700, scan rate 10 Hz, ID disabled) for the two most intense ions per scan. The ions were selected when they reach an intensity threshold of 1500, isolated at the tuning file Q1 resolution, fragmented using collision induced dissociation (CID) with fixed collision energy (CE 20 eV), and excluded for 1 s before being re-selected for fragmentation. For the ESI source, the parameters were set to interface voltage 4 kV, interface temperature 300 °C, nebulizing gas flow 3 L/min, and drying gas flow 10 L/min. The parameters used for the CDS probe include an interface voltage 4.5 kV, and nebulizing gas flow 1 L/min.

### Comparative metabolomics

Raw LC-MS data were converted to open source mzXML format using LabSolutions software (Shimadzu), and the converted files were imported into MZmine 3.3.0^59^ for data processing. Unless specified otherwise, *m/z* tolerance was set to 0.002 *m/z* or 10.0 ppm, RT tolerance was set to 0.05 min, MS1 noise level was set to 1.0E3, MS2 noise level to 1.0E1 and the minimum absolute height was set to 5.0E2. The option to detect isotope signals below noise level was selected. For feature detection and chromatogram building, the ADAP chromatogram builder^60^ was used with positive polarity, centroid mass detector, minimum group size of 5 in number of scans and a 2.0E3 group intensity threshold. The obtained peaks were smoothed (width: 9), and the chromatograms were deconvoluted using the local minimum search with a 90% chromatographic threshold, 1% minimum relative height, minimum ratio of peak top/edge of 2 and peak duration of 0.03 to 3.00 min. The detected peaks were deisotoped (monotonic shape, maximum charge: 5; representative isotope: most intense). Peak lists from different extracts were aligned (weight for *m/*z: 20, weight for RT: 20, compare isotopic pattern with a minimum score of 50%). The gap filling algorithm was used to detect and fill missing peaks (intensity threshold 1%, RT tolerance: 0.1 minute). Duplicate peaks were filtered, and artifacts caused by detector ringing were removed (*m/z* tolerance: 1.0 m/z or 1,000.0 ppm). The aligned peaks were exported to a MetaboAnalyst. From here, peaks were additionally filtered to keep only peaks present in all 3 replicates and not in the media blanks, using in-house scripts. The resulting MetaboAnalyst peak list was uploaded to MetaboAnalyst^61^, log transformed, and normalized with Pareto scaling without prior filtering. Missing values were filled with half of the minimum positive value in the original data. Volcano plots were generated using default parameters. Additionally, extracted ion chromatograms have been obtained for the ions of the DFO-related metabolites (*m/z* tolerance 0.001 or 5 ppm, Table S4). An in-house python script was used to visualize these chromatograms with matplotlib v3.7.2 pyplot^62^.

### Plasmids, constructs and oligonucleotides

All plasmids and constructs described in this work are summarized in Table S5. The oligonucleotides are listed in Table S6.

Fragment containing *gapdh* promoter was digested from previously published plasmid pGWS1370^63^ and cloned into pCRISPR-cBEST^34^ via the same restriction sites to generated pGWS1384, where the expression of Cas9n (D10A), cytidine deaminase and uracil-DNA glycosylase inhibitor (UGI) were under the control of *gapdh* promoter instead of *tipA* promoter. Spacers of each targeted gene were selected on CRISPy-web^64^ and cloned into NcoI-digested pGWS1384 via single strand DNA (ssDNA) oligo bridging method. Single strand DNA (ssDNA) oligos SCO4048_W55 and SCO4048_Q68b were used to generate SCO4048 knockout constructs pGWS1582 and pGWS1584, respectively. Similarly, SCO4049 knockout construct pGWS1585 was created using oligo SCO4049_W61. SCO4050 knockout constructs pGWS1598 and pGWS1590 were created employing oligos SCO4050_W43 and SCO4050_Q91, respectively. All the generated knockout constructs were validated by Sanger sequencing using primer sg_T7_R_SnaBI.

For the complementation of SCO4048 null mutant, pGWS1596 was used, an integrative vector based on pSET152 and harboring SCO4048 under the control of *gap* promoter. The *gap* promoter and the entire coding region (+1/+724) of SCO4048 were amplified from *S. coelicolor* M145 genomic DNA using primer pairs Pgap_F and Pgap_R, and SO4048_F and SCO4048_R, respectively. Fragments were cloned into EcoRI and XbaI digested pSET152 via Gibson assembly to generate pGWS1596. Similarly, pGWS1597 and pGWS1598 were created for the complementation of SCO4049 and SCO4050 null mutants, respectively. The coding region (+1/+2347) of SCO4049 in pGWS1597 was amplified using primers SCO4049_F and SCO4049_R, while the coding region (+1/+619) of SCO4050 in pGWS1598 was amplified using primer pair SCO4050_F and SCO4050_R.

## Supporting information

All Supplemental Information

## ACKNOWLEDGEMENTS

The work was supported by the European Union via ERC Advanced Grant 101055020-COMMUNITY to G.P.v.W. and ERC Starting Grant 948770-DECIPHER to M.H.M.

