## Supplementary material for "Prediction of gene cluster function based on transcriptional regulatory networks uncovers a novel locus required for desferrioxamine B biosynthesis": All Supplemental Information

Belonging to the manuscript

#### Supplementary Figures

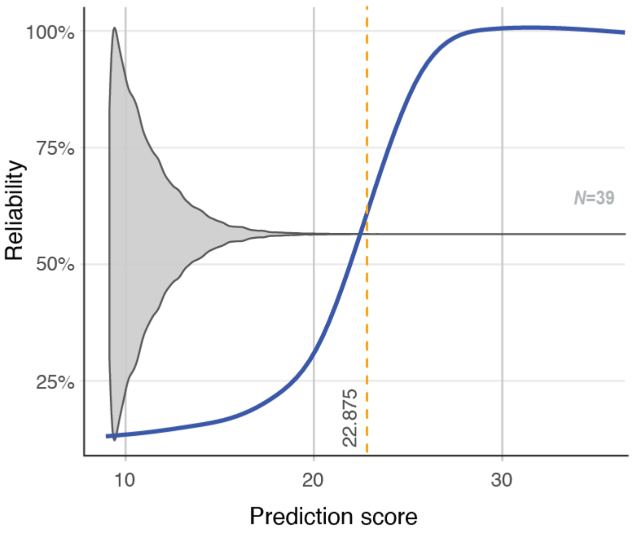


**Figure S1.** DmdR1 PWM threshold setting and identification of novel iron boxes in *S. coelicolor*. Gray violin plot of the distribution of the number of matches to the PWM. The blue S-curve indicates the ratio (in %) of hits in the non-coding versus coding regions of the genome. The orange dotted line is the threshold set by the median score of the hits.

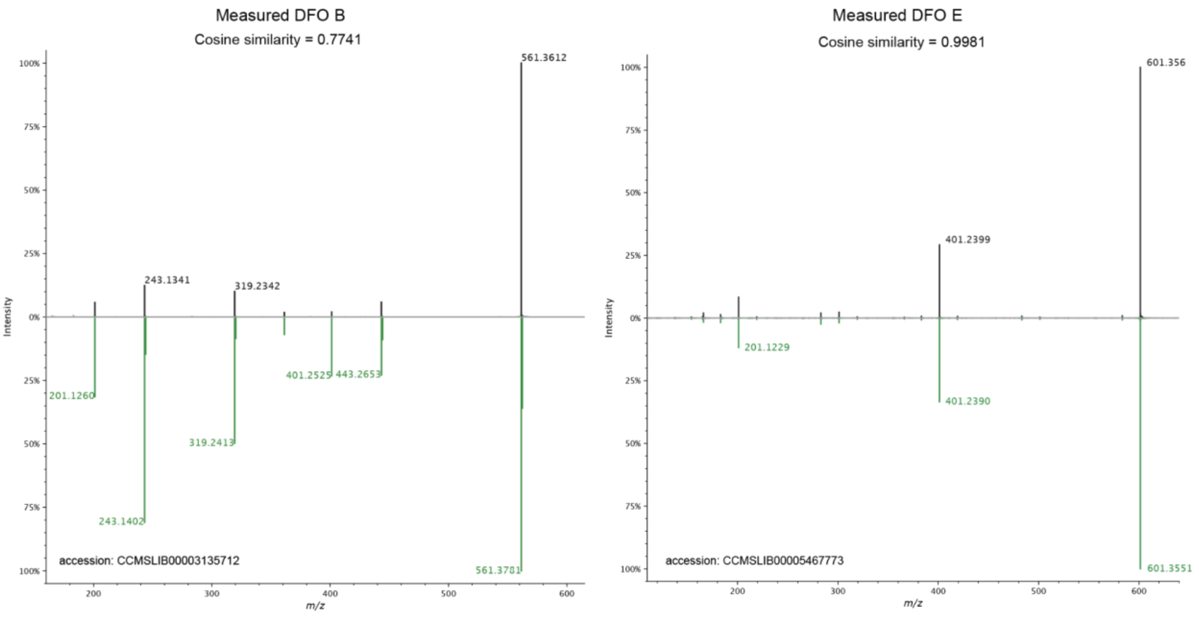

**Figure S2.** Measured MS and MS/MS spectra of desferrioxamine (DFO) B and E compared to their respective GNPS library spectra with accessions CCMSLIB00001059066 and CCMSLIB00005467773 of GNPS Mirror Match [1–4].

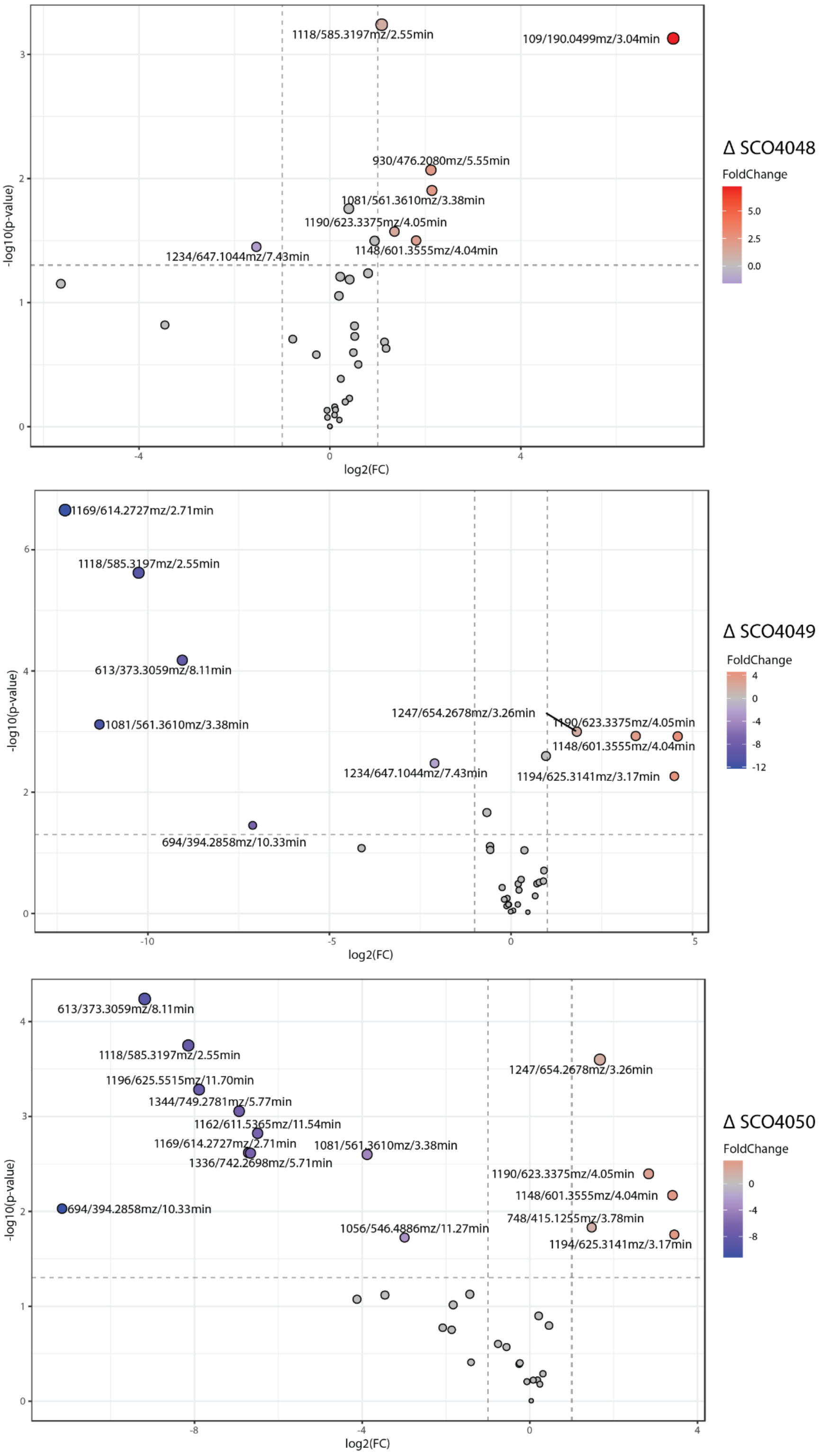

**Figure S3.** Volcano plots showing the relative intensities of the mass features detected in the generated knock-out mutants of *S. coelicolor* in comparison with the wild-type. Mass features were regarded as significantly upregulated (red colored) or downregulated (blue colored) in the knock-out strains, with at least a two-fold change in intensity and $p$ value ≤ 0.05.

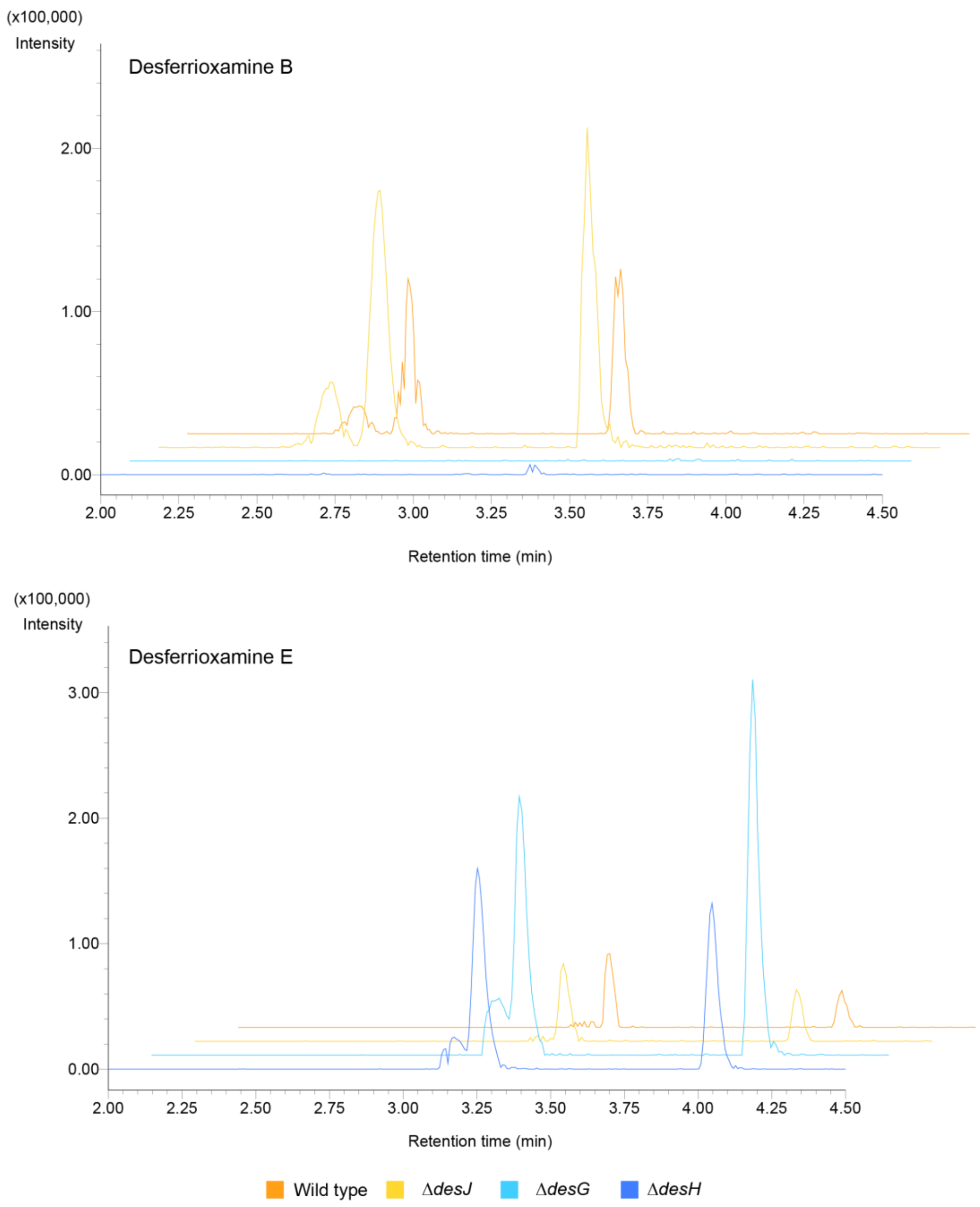


**Figure S4.** Extracted ion chromatograms of DFOB and DFOE related metabolites in the knock-out (Δ) mutants of SCO4048 (*desJ*), SCO4049 (*desG*) and SCO40450 (*desH*) compared to the parent *S. coelicolor* M145 strain.

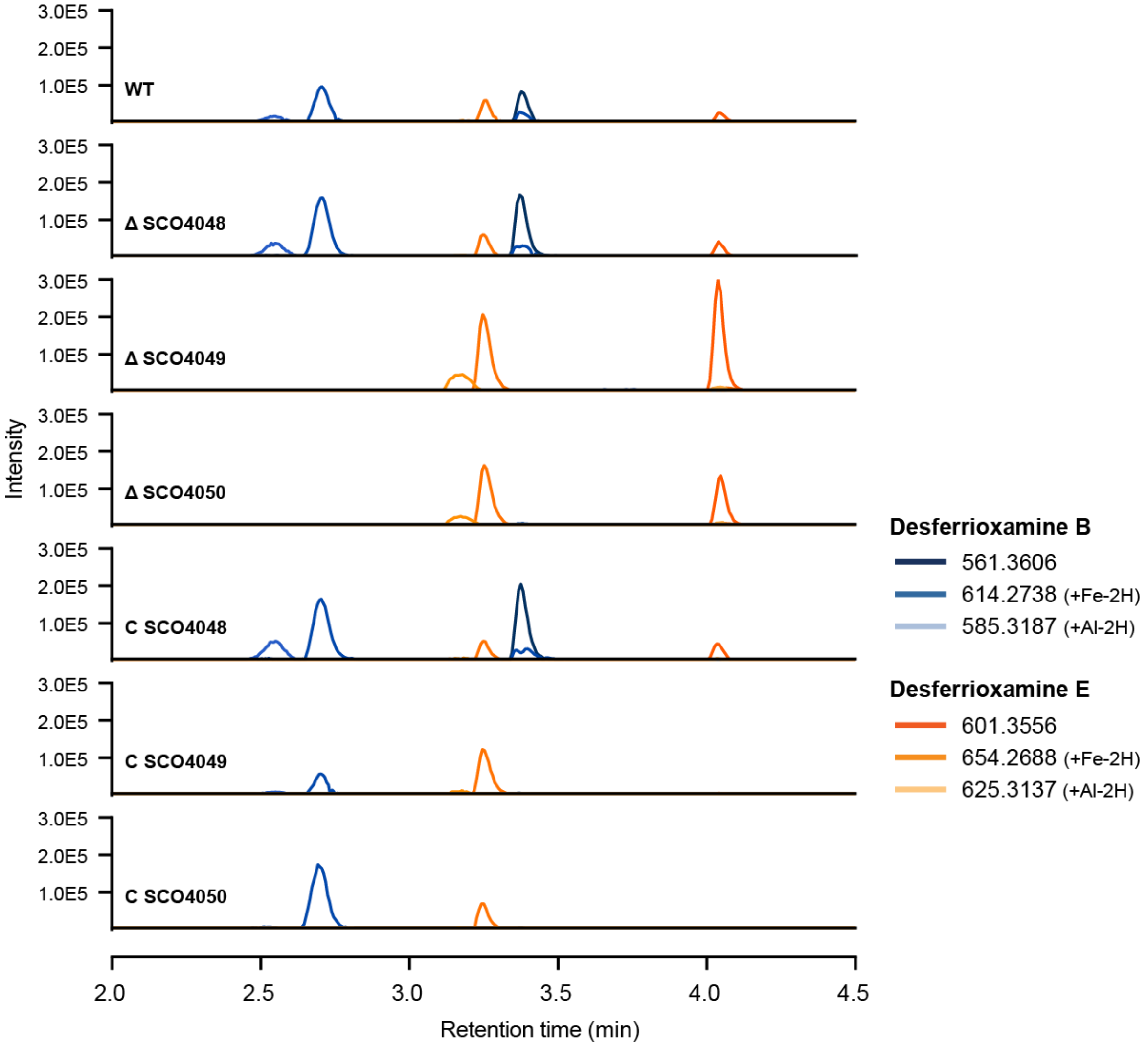


**Figure S5.** Extracted ion chromatograms of DFO-related metabolites in the knock-out (Δ) and complementation (C) mutants of SCO4048 (*desJ*), SCO4049 (*desG*) and SCO40450 (*desH*) compared to the parent *S. coelicolor* M145 strain.

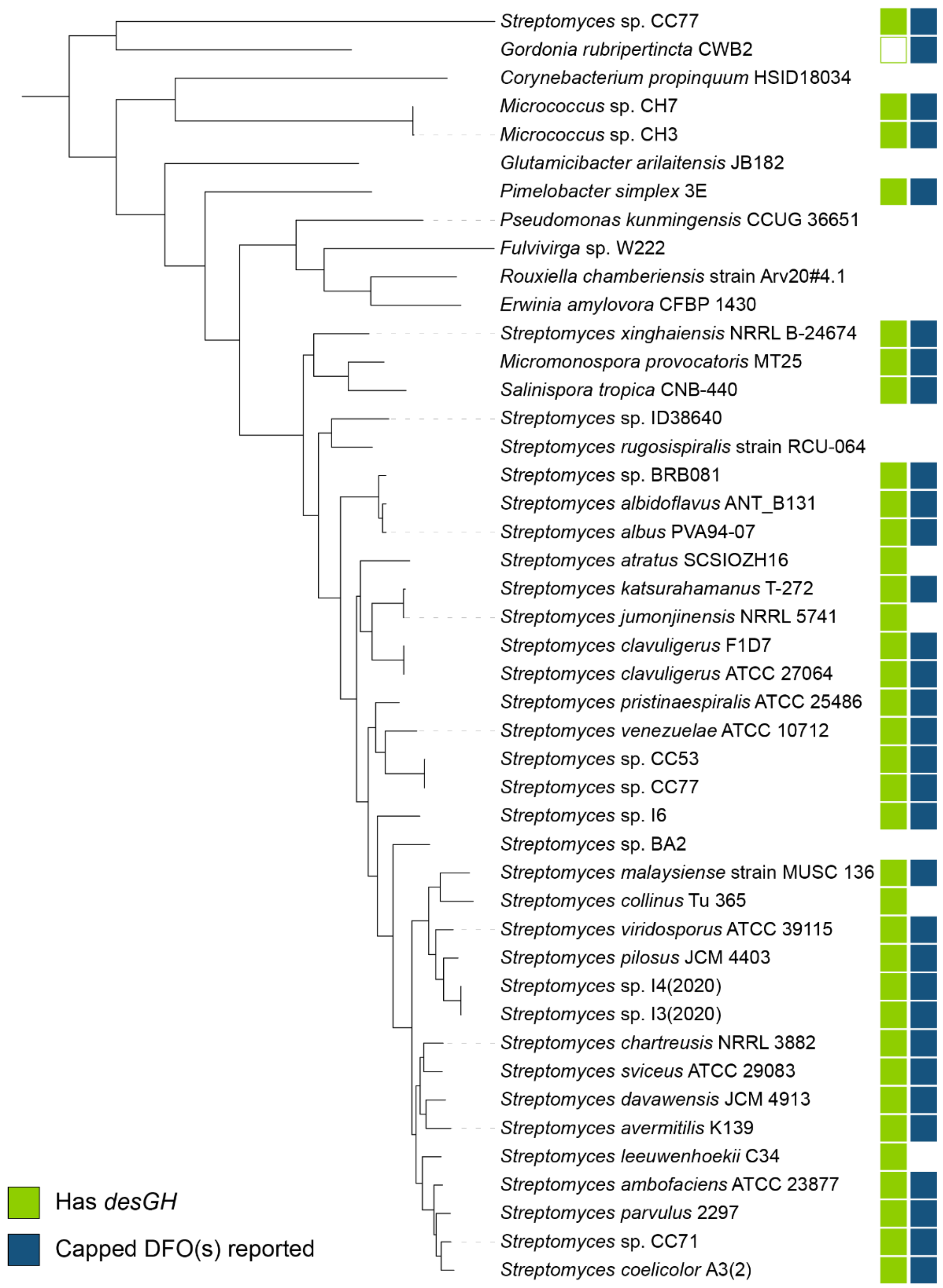


**Figure S6.** Midpoint-rooted phylogenetic gene tree of the *desABCD* locus in previously reported DFO producers (Table S3), produced by CORASON.[5] Green squares indicate that the genome of the strain also contains homologs of *desGH* in addition to *desABCD*, and dark blue squares indicate that the strain was reported to produce acetyl, fatty-acyl, or aryl “capped” DFOs. The genome of *Gordonia rubripertincta* CWB2 contains a homolog of *desG* in an expanded locus (see SI discussion 2).

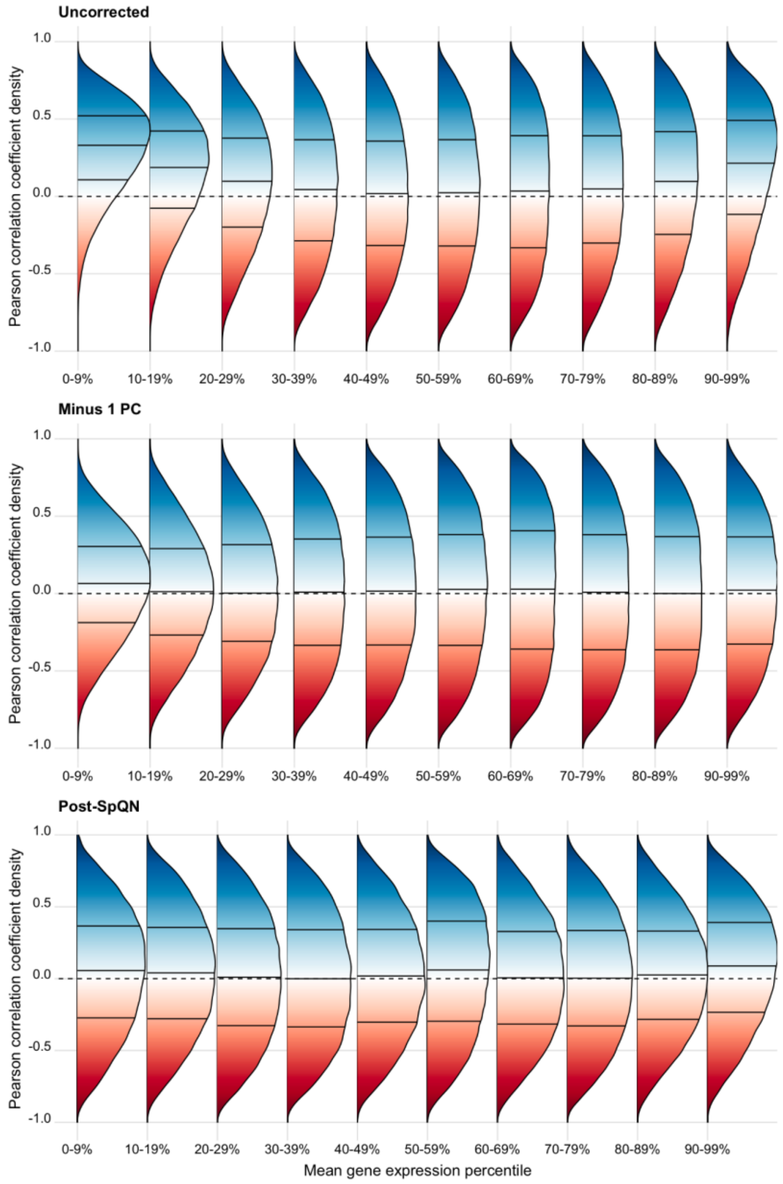


**Figure S7.** Mean-expression biases in correlation coefficient distributions after standard processing (top), after regression of the first principal component (middle), and after spatial quantile normalization (bottom). Genes were sorted by mean expression and split into 10 bins of equal size. All-to-all Pearson correlation coefficients were calculated within each bin, and the density was calculated and plotted with the R package *ggridges*. Deviations from a zero-centered distribution (dashed line) suggest non-biological confounders. Solid black lines give the first, second, and third quartiles.

### Supplementary discussion

#### SI Discussion 1: Description of the putative DmdR1 regulon in *S. coelicolor*

DmdR1 uses Fe(II) as a co-repressor, and thus the intracellular concentrations of both iron and DmdR1 will affect the expression of the regulon. Although the majority of research on metal-sensitive regulatory proteins focuses on the metal concentration, the anti-correlation in the expression between *dmdR1* and its putative target genes (Figure 2) suggests a significant role for DmdR1 concentration under the conditions tested by Lee *et al*. Most of the target genes are more tightly co-expressed with each other (median PCC = 0.90) than anti-correlated with *dmdR1* expression itself (median PCC = –0.66, $p$ = 1.2e-8, one-sided Mann-Whitney U test). Intracellular Fe(II) concentrations or other post-transcriptional factors likely decouple the expression of *dmdR1* and its regulon. At least two other repressors control the expression of *dmdR1*; these are DasR (in the absence of glucosamine-6-phosphate),[6] and the phosphate master regulator PhoP (during phosphate starvation).[7] In the processed RNA-Seq dataset, the expression of *dmdR1* and *dasR* did not correlate (PCC = -0.02), while *dmdR1* and *phoP* showed anti-correlated transcription (PCC = -0.71). However, aspects of post-transcriptional control or not considered, such as by nutrients or posttranslational modification. For example, the DNA binding activity of DasR is controlled by different phosphates, which connects the PhoP and DasR regulons[39].

##### *Siderophore-mediated iron acquisition*

The majority of the DmdR1 regulon is putatively involved in the control of siderophore pathways. In total, we identified four DmdR1 binding sites in the coelichelin locus (*cch*, **SCO0489-0499**) and two binding sites in the desferrioxamine locus (*desABCD*, **SCO2780-2785**), as well as one binding site upstream of *desGH* (**SCO4049-4050**), as discussed in the main text. After biosynthesis, the ABC transporter complex CchGI (SCO0491 and SCO0493) is presumed to export coelichelin from the cell.[8] The desferrioxamine locus does not encode a putative export protein; one of the DmdR1-controlled major facilitator superfamily (MFS) transporters encoded by **SCO0680** or **SCO6822** may be responsible. Iron-bound siderophores are transported back into the cell through ABC transporter complexes. The coelichelin locus encodes one complete uptake system, CchCDEF (**SCO0494-0497**), while the desferrioxamine binding protein DesE (**SCO2780**) is proposed to partner with the incomplete ABC complex encoded by **SCO1785-1787**. A third, standalone iron-siderophore ABC importer has also been characterized, CdtABC (**SCO7400-7398**).[9,10] **SCO7397,** putatively in operon with *cdtABC*, encodes part of the RNase T / DNA polymerase III family of exonucleases (IPR013520); any relation to iron-starvation is unclear. **SCO6426** shows homology to an ABC transporter ATP-binding component; however, ATP-binding residues found in homologues are absent.[11] SCO6426 may be the remnant of an ancestral siderophore ABC transporter system which can still be found complete in other *Streptomyces* species (*e.g.* in *S. olivaceus*, KVH11_RS17020-35). After entering the cell, siderophore-bound Fe(III) is reduced by a "siderophore-interacting protein" (SIP) that allows for iron release.[12] The desferrioxamine locus encodes one SIP reductase, DesF (**SCO2781**),[8] while putative SIPs **SCO0459** and **SCO1623** may be involved in iron release from coelichelin and/or xenosiderophores. A fourth SIP DesJ (**SCO4048**) may encode a desferrioxamine B-specific reductase, as evidenced by increased levels of desferrioxamine B in the supernatant of a *desJ* mutant (main text).

##### *Siderophore-independent iron acquisition*

**SCO2267** to **SCO2277** are eleven co-expressed genes that contain three predicted DmdR1 binding sites. None of these genes have been characterized, but they are homologous to siderophore-independent iron uptake systems in other organisms. **SCO2275-2277** are homologous to *efeO*, *efeB*, and *efeU*, respectively, which are involved in an *E. coli* transport system that imports Fe(II) under aerobic, low-pH conditions.[13] A homologue of **SCO2268** in *Bordetella bronchiseptica* (BB3589) is adjacent to an unrelated low-pH ferrous iron transport system, *ftrABCD*.[14] However, a BB3589 null mutant showed no growth defect, and likewise the exact function of SCO2268 is unclear. **SCO2267** and **SCO2270-2274** are homologous to *Corynebacterium diptheriae* genes *hmuO*, *htaA*, and *hmuTUV* that are required for acquiring iron from heme and hemoglobin.[15 16 17,18] Distant gene **SCO0799** encodes a HugZ-like heme oxygenase that may also be involved in Fe-heme utilization.[19,20] **SCO2269** is a hypothetical protein that does not match any known protein families; however, its location suggests a role in a siderophore-independent iron acquisition pathway.

##### *Mobilization of stored iron*

Under iron-replete conditions, *S. coelicolor* stores excess iron as a ferric oxide biomineral within a bacterioferritin nanocage, encoded by SCO2113.[21] Neighbouring gene **SCO2114**, which encodes a bacterioferritin-associated ferredoxin (Bfd), is associated with a DmdR1 binding site and is therefore likely controlled by it. This family of ferredoxins mobilizes stored iron for metabolic use by transferring electrons to bacterioferritin for reductive demineralization.[21–23] In turn, **SCO0681** encodes a ferredoxin reductase, which could serve as the terminal electron source for Bfd.

##### *Links to other pathways*

Unchelated intracellular iron catalyzes the formation of reactive oxygen species, linking iron acquisition to oxidative stress responses.[24] **SCO5830** encodes a putative thioredoxin-like ferredoxin that was previously found to be highly expressed under copper overload conditions,[25] and thus may be involved in a shared response to metal-induced reactive oxygen species. **SCO2350** is in the DoxX-like family of membrane proteins, members of which are upregulated during oxidative stress in *Mycobacterium tuberculosis* and *Haemophilus influenzae*.[26,27]

**SCO6159**, a GntR-type regulator of the FadR sub-family,[28] was the only gene downstream of a putative DmdR1 binding site that was strongly co-expressed with DmdR1 itself (PCC = 0.76). The divergently-transcribed **SCO6158** is not a match to any known protein families except for a short C-terminal domain with a [2Fe-2S] binding motif that is identical to that of the siderophore reductase FhuF.[29] This iron-sulfur cluster may be involved in electron transport, like in FhuF itself, or in sensing environmental signals for a regulatory pathway.[30]

The presence of a DmdR1 binding site in the intergenic region between SCO5998 and SCO5999 links DmdR1 to primary metabolism. **SCO5998** (MurA2) encodes one of two uridine diphosphate N-acetylglucosamine (UDP-GlcNAc) transferases in *S. coelicolor*, which catalyze the first committed step of peptidoglycan biosynthesis for cell walls.[31] A DasR-mediated link between GlcNAc and DmdR1 has previously been established.[6] **SCO5999** encodes aconitase (AcnA), which isomerizes citrate to isocitrate in the citric acid cycle.[32] A secondary role for AcnA has been reported in *E. coli* and *B. subtilis*, and proposed in *S. coelicolor*: under iron starvation or oxidative stress, a [4Fe-4S] cluster disassembles, and AcnA becomes a post-transcriptional regulator of gene expression that binds to mRNA sequences called “iron responsive elements”.[32,33] While SCO5998 expression was anti-correlated with *dmdR1*, SCO5999 expression was uncorrelated.

The SCO4046-4052 cluster contains *desGHJ*. The other four genes have no obvious relationship to iron homeostasis. The hypothetical protein encoded by **SCO4046** is predicted to be intrinsically disordered by MobiDB.[34] The predicted gene product of **SCO4047** is homologous to the HhH-GPD superfamily of base excision repair enzymes.[35] **SCO4051** and **SCO4052** encode a putative sugar epimerase and dehydrogenase, respectively.

#### SI Discussion 2: Identification of a putative nocardichelin biosynthetic gene cluster

*Gordonia rubripertincta* CWB2 was found to produce acetyl-capped desferrioxamines B and A1, but the genome contains only a *desG* homolog without *desH*.[36] A manual inspection of the biosynthetic gene cluster reveals a distinct GCN5-related acetyltransferase GCWB2_02925 that may be responsible for the acetylated desferrioxamines. The *G. rubripertincta* CWB2 siderophore locus also encodes for mycobactin-like salicylate synthase and non-ribosomal peptide synthetase (NRPS) genes (Table S7), all of which were upregulated under low-iron conditions.[36] Only one NIS/NRPS hybrid siderophore family has been identified to date, nocardichelins A and B from the unsequenced *Nocardia* strain Acta 3026.[37] The *G. rubripertincta* CWB2 gene cluster is consistent with the biosynthesis of nocardichelins, as proposed in Figure S7. Furthermore, nocardichelins were recently detected in extracts from unsequenced *Gordonia* strain Berg02-22.2.[38]

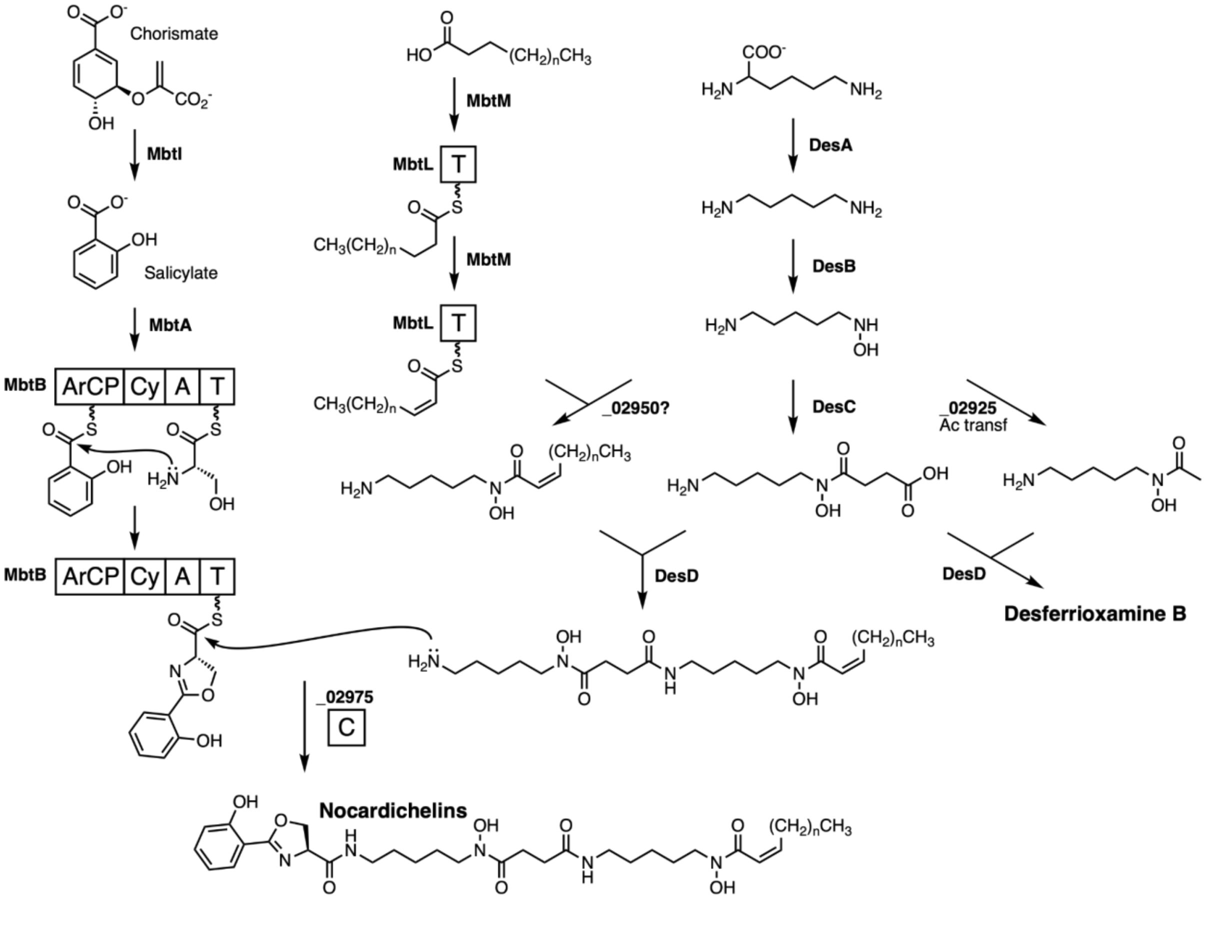


**Figure S8.** Proposed biosynthesis for nocardichelins and desferrioxamine B in *Gordonia rubripertincta* CWB2. Gene names correspond to Table S7, while underscores indicate the locus tag prefix “GCWB2_”.
